# GABA-induced Ca^2+^ signaling in the primary cilium of neurons

**DOI:** 10.1101/2025.05.26.656109

**Authors:** Gonzalo Sanchez, Moa Södergren, Devendra Kumar Maurya, Alfhild Grönbladh, Maria Lindskog, Olof Idevall-Hagren

**Affiliations:** Department of Medical Cell Biology, Uppsala University, Uppsala, Sweden; Department of Molecular Biology, Umeå University, Umeå, Sweden; Department of Pharmaceutical Biosciences, Uppsala University, Uppsala, Sweden

**Keywords:** GABA, GABA-B1, Ca^2+^, cilium, neuronal compartments

## Abstract

The complex signaling processing in neurons requires establishment of autonomous compartments. The compartment that is unique for neurons and crucial for neuron-neuron signaling is the synapse. Another compartment, that neurons share with all other cells in the body, is the primary cilium. The primary cilium is a solitary organelle, present in almost every neuronal cell type, that extends into the extracellular space for detection of signals. Several GPCRs have been identified as ciliary receptors, and here we show that the metabotropic GABA receptor subtype 1 localizes to primary cilia of neurons across different regions of the mouse brain and that activation of these receptors initiates Ca^2+^ signaling that is restricted to this organelle. The excitatory nature of GABAergic signaling in primary cilia is opposite to GABA action in other neuronal domains, indicating distinct modes of action of this universal inhibitory neurotransmitter even within the same neuron.

## INTRODUCTION

The primary cilium is a slender appendage protruding from the apical membrane of most cell types. It has a conserved ultrastructure consisting of nine pairs of microtubules doublets which provides mechanical support and a route for bidirectional transport of particles along the cilium. It is the only organelle that is in contact with the extracellular milieu and its primary function is to sense changes and to prompt appropriate cellular responses, yet fundamental understanding of how this sensing is accomplished and how signals from the cilium are transduced to the soma is largely missing. The best-understood ciliary signaling pathway is that of the developmental morphogen Hedgehog (Hh) ^1^. Binding of Hh to its receptor Patched results in the removal of the receptor from the cilium which in turn enables Smoothened, an orphan GPCR, to enter the ciliary compartment and promote activation of GLI transcription factors ^2^ ^3^. Many other GPCRs, including those of several neurotransmitters and neuromodulators such as serotonin, dopamine, and somatostatin also localize to the cilium ^4^ ^5^ ^6^, but the importance of this localization is not clear, in particular since many of these receptors also localize to other cellular domains. Recently, it was shown that neuronal primary cilia are positioned in proximity to synapses where they may detect synaptic spillover, and certain neurons can even form direct synapses with primary cilia ^7^ ^5^ ^8^. Activation of ciliary receptors initiates local formation of second messenger that, through largely unknown mechanisms, regulates diverse neuronal functions, including control of gene expression and of excitatory to inhibitory balance within neuronal circuits ^4^ ^9^ ^10^.

Ca^2+^ is a key second messenger in neurons and Ca^2+^ signals are both compartmentalized and dynamic. Ca^2+^ signaling can be restricted to different organellar compartments, but also to compartments without physical boundaries, such as dendritic spines and nerve terminals ^11^ ^12^. In the latter case, local Ca^2+^ signals are maintained through the concerted action of local influx and extrusion that, together with the high Ca^2+^ buffering capacity of cells, generate Ca^2+^ microdomains. The primary cilium, similar to a dendritic spine, is not fully enclosed by membranes, but is connected to the cytosol through an opening at its base. We recently showed that primary cilia in insulin-secreting cells constitute a unique Ca^2+^ compartment that is isolated from the influence of cytosolic Ca^2+^ as a consequence of efficient extrusion at the cilia base ^13^. This enables cilia to initiate intrinsic Ca^2+^ signaling downstream of cilia-localized receptors without cytosolic interference. If similar principles govern Ca^2+^ signaling in neuronal primary cilia is not known.

GABA is the main inhibitory neurotransmitter in the adult mammalian brain where it acts by engaging either ionotropic GABA-A or metabotropic GABA-B receptors. GABA-A receptors are chloride channels that can drive hyperpolarization and shunting for damping neuronal excitability ^14^. They have been found in almost all cellular domains with the highest enrichment at postsynaptic sites opposite to release terminals but also at extra-synaptic dendritic and somatic membranes as well as the axon initial segment. By exerting their inhibition at specific subcellular locations, they can influence neuronal computational capabilities in distinct ways. GABA-B receptors couple to G-proteins and regulate K^+^ and Ca^2+^ channels at both pre- and post-synaptic locations ^15^. In addition, we recently found that GABA-B1 receptors were strongly enriched in primary cilia of hormone-secreting cells in the pancreas, that this localization was modulated by GABA binding and that receptor activation triggered ciliary Ca^2+^ signaling ^13^. This paradoxical excitatory effect of GABA involved local voltage-dependent influx of Ca^2+^, but whether this mechanism exists outside of the endocrine pancreas is not known. In particular, it would be highly relevant to determine if GABA has previously unknown signaling capabilities in the brain, and whether these might involve primary cilia.

We now show that GABA-B1 receptors localize to the base of primary cilia in neurons across different brain regions, but are absent from the cilia of non-neuronal cells. The receptor localized to the cilia membrane and the density in the cilium was even higher than in adjacent dendrites. Direct measurements of ciliary Ca^2+^ concentration changes in cultured neurons revealed that primary cilia were largely isolated against Ca^2+^ concentration changes in the soma, and that activation of GABA-B1 receptors triggered Ca^2+^ signaling that was restricted to the cilia.

## METHODS

### Materials

5HT6-G-GECO (Addgene plasmid 47499; gift from Takanari Inoue, Johns Hopkins University, Baltimore, MD) ^16^ was packaged into a serotype E5 adenovirus (Vector Biolabs) and used to transduce neuronal cultures. The following primary antibodies (1/300 dilutions) were used: ARL13b (Abcam ab136648), AC3 (Abcam ab277619), AC3 (Alomone AAR-043), NeuN (Sigma-Aldrich ABN90), GFAP (Synaptic System 173–004), Synapsin II (Alomone ANR-015), GABA-B1 (Alomone AGB-001), MCHR1 (Thermo Fischer PA5-77492), Patched (Abcam ab53715), EP4 (Santa Cruz Biotechnology sc-55596), ANKS6 (Sigma HPA008355), NPHP3 (Proteintech 22026-1-AP), NEK8 (kind gift from Prof. David R. Beier, Seattle Children’s Hospital, USA). The following secondary antibodies (diluted 1/500) were used for confocal microscopy: Anti-mouse 568 nm (A11004; Invitrogen) and anti-mouse 488 nm (A28175; Invitrogen), anti-mouse 647 nm (A31571), anti-rabbit 488 nm (A28175; Invitrogen), anti-rabbit 568 nm (A11011), anti-guinea pig 568 nm (A11075) and anti-guinea pig 647 nm (A21450). The following secondary antibodies were used for STED microscopy: Star 580 (Abberior ST580-1001) and Star Red (Abberior STRED-1002).

### Animals

All animal experiments were in accordance with Swedish rules and guidelines for animal experiments (Animal Welfare Act SFS: 2018:1192) and the European Union directive on the Protection of Animals Used for Scientific Purposes (Directive 2010/63/EU). The animal experiment protocol was approved by the local animal ethics committee in Uppsala.

### Animal perfusion

6-months-old C57BL/6 wild type mice (Charles River, Germany) were anesthetized with a solution of ketamine (100 mg/kg) and xylazine (16 mg/kg) i.p. After disappearance of corneal reflexes, the mice were transcardially perfused first with NaCl 0.9%, followed by PFA 4%. The brains were extracted and fixed in PFA 4% for 48h, thereafter washed in PBS for 24h and submerged in a solution of 30% sucrose and 1% sodium azide until further use.

### Olfactory and respiratory tissue preparations

Nasal regions of one month old postnatal C57BL/6 WT mice were fixed in 4% paraformaldehyde at 4°C for five hours and decalcified in 0.5 M EDTA, pH 8.0, for 48 hours at 4°C. Decalcified tissues were incubated in 30% sucrose (w/v in PBS) overnight and frozen in OCT (Histolab, catalogue # 45830). The OCT blocks were sectioned using an HM 550 Cryostat Microtome at a thickness of 14 µm. Air-dried sections were incubated with citrate buffer (10 mM, pH 6.0) for 5 min at 100 °C, washed in PBS and incubated in blocking solution (2% FCS + 0.2% Triton X-100 in PBS). The sections were washed in PBS, incubated overnight with primary antibodies (OMP, catalogue # 019-22291, Fujifilm wako chemicals. CNGA2, catalogue # sc-1370, Santa Cruz Biotech Inc. Acetylated tubulin, catalogue # T7451, clone 6–11B-1 Sigma. GABA(B)R1, catalogue # AGB-001, Alomone labs) in blocking solution, washed and incubated with secondary antibodies (Dylight 488-conjugated anti-mouse, catalogue # AS101201, Agrisera AB. Alexa 546-conjugated anti-rabbit, catalogue # A-10040, Life Technologies. Alexa 488-conjugated anti-goat, catalogue # A-11055, Life Technologies) diluted in T-PBS (0.1% Tween-20 in PBS). The sections were then stained with Hoechst (0.1 µg/ml) and mounted with fluorescence mounting media (Dako, catalogue # S302380-2). Imaging was performed with a Leica TCS SP8 confocal system equipped with a Leica DMi8 microscope using an HC PL APO CS2 63x/1.40 N.A. objective. Confocal images with a z-axis were acquired using LAS X software.

### Neuronal culture

Foetuses of pregnant Sprague Dawley dams (Charles River, Italy) were used to set up primary cortical and hippocampal cell cultures. The rat primary cell cultures were collected from embryonic day 17 Sprague Dawley foetuses by harvesting part of the cortex and the hippocampus, as described earlier ^17^. Briefly, the tissue was digested and dissociated into a homogenous cell suspension and dissolved in Gibcos neurobasal plus media (NBM; Thermo Fisher Scientific, Waltham, MA, USA) supplemented with 0.25% glutaMAX™ (Thermo Fisher Scientific), 1% penicillin/streptomycin (Thermo Fisher Scientific), and 4% Gibcos B27 plus (Thermo Fisher Scientific). Primary cultures were grown on poly-D-lysine coated #1 glass coverslips with cell density around 150.000 cells/cm^2^, and kept in serum-free Neurobasal medium supplemented with 0.5 mM GlutaMAX and 4% (v/v) B27. Cultures were maintained at 37°C and 5% (v/v) CO_2_ in a humidified incubator and partial media changes were performed twice a week. Cultures were kept for no less than three weeks before the viral-induced expression of ciliary calcium reporters, followed by another two days at least for allowing sufficient expression.

### Mouse Islets

Adult C57BL/6J mice (>8 months old) were sacrificed by CO_2_ asphyxiation and decollation, and the pancreas was removed and put on ice prior to digestion with collagenase P and mechanical disaggregation at 37°C. The digestion was terminated by 1 ml BSA 0.1 g/ml, islets were then separated from exocrine tissue by hand-picking with a 10 µl pipette under a stereo microscope. Islets were cultured in RPMI 1640 medium with 5.5 mM glucose, supplemented with 10% fetal calf serum, 100 units/ml penicillin, 100 µg/ml streptomycin for 2–5 d at 37°C in a humidified atmosphere with 5% CO_2_.

### MIN6 pseudoislets

Detached mouse insulinoma β-cells (MIN6 cells; 3 to 5 million/ml passages 18–30) ^18^ were transferred to a non-adherent Petri dish (Sarstedt) with 5 ml culture medium, put onto low-speed orbital shaker and kept in culture for 5–7 d at 37°C in a humidified atmosphere with 5% CO_2_ until they spontaneously formed cellular aggregates (pseudoislets). The culture medium was based on DMEM (Life Technologies) supplemented with 25 mmol/liter glucose, 15% FBS, 2 mmol/l L-glutamine, 50 μmol/l 2-mercaptoethanol, 100 U/ml penicillin, and 100 μg/ml streptomycin.

### Immunohistochemistry

Mouse brains were cut using a cryostat into 40 µm thick slices which were stored in antifreeze solution containing 0.05 M sodium phosphate buffer, 0.44 M sucrose and 30% ethylene glycol (pH 7.6) at -20°C until usage. The brain slices were first subjected to antigen retrieval (Tris 10 mM EDTA, 10% Tween-20) at 80°C for 30 minutes. Sections were blocked and permeabilized using 10% normal donkey serum (NDS) and 0.3% Triton X-100 in PBS for one hour at room temperature, and subsequently incubated with primary antibodies in same blocking solution for 72 hs. Sections were then washed and incubated with Alexa Fluor-conjugated secondary antibodies (Invitrogen, Waltham, MA, USA) in blocking solution for two hours. After washing, sections were mounted in mounting medium (Fluoromount, Sigma Aldrich, St. Louis, MO, USA) on glass slides and covered with clean glass coverslips (#1 for confocal and #1,5 for STED microscopy).

### Immunocytochemistry

Neuronal primary cultures were fixed with 4% paraformaldehyde for 10 minutes followed by permeabilized with 0.3% Triton-X (diluted in D-PBS) for 20 minutes and blocking for 30 minutes in same solution supplemented with 5% bovine serum at room temperature. Incubation with primary antibodies was performed overnight at 4^°^C. The next day coverslips were washed with D-PBS and then incubated with the secondary antibodies for 1 hour in blocking solution. Finally, coverslips were rinsed with D-PBS and mounted with ProLong Gold Antifade reagent.

### Viral transduction

After no less than 3 weeks following plating of both cortical and hippocampal primary neurons, culture medium was removed from the 6-well-plates where coverslips were kept and replaced with 0,6 ml of culture medium containing the adenovirus (5 µl of high titration virus, >10^12^ - 10^13^ vp/ml diluted in 7,2 mL of culture medium). After 3 h of incubation, the virus containing medium was replaced by fresh medium (2 ml/well) and the culture kept for at least 48 h to allow for biosensor expression.

### Ciliary Ca^2+^ imaging

Recordings were performed using a Nikon Eclipse Ti2 microscope body equipped with a 100x 1,49NA oil objective, a Yokogawa CSU-X1 spinning disk confocal head with a 483/32 nm emission filter and an Andor iXon-EMCCD camera. Single wavelength imaging was performed using 491nm Cobolt Calypso laser with an acquisition frequency set at 1 picture per 0,5 second. In a subset of experiments the calcium reporter dye Cal590 was included in order to improve somatic calcium detection in a separate channel with illumination relying on 568nm Cobolt Jive laser and a 600/52 emission filter. Image acquisition was controlled via Metamorph software running on a PC with Windows 10 operating system. Experiments were performed in an imaging buffer containing the following salts (in mM): 125 NaCl, 4.9 KCl, 1.3 MgCl_2_, 1.2 CaCl_2_ and supplemented with (in mM) 10 glucose, 25 Hepes and 1 mg/ml BSA with pH set to 7,4 and all recordings were done at room temperature and without perfusion; all drugs were directly added to the buffer in the recording chamber at the adequate concentration to achieve desired working dilutions. A few recordings were performed in TIRF configuration as described in Sanchez et al., 2023, and since Ca^2+^ dynamics did not differ with that observed in confocal configuration data was pooled together (spontaneous calcium activity). Briefly, TIRF imaging was performed on a Nikon TiE microscope equipped with an iLAS2 TIRF illuminator for multi-angle patterned illumination (Gataca systems) and a 60x 1.49NA Apo-TIRF oil-immersion objective. Excitation light for GGECO delivered by 488 nm diode-pumped solid-state laser with built-in acousto-optical modulators (all from Coherent, Inc). Fluorescence was detected with a back-illuminated EMCCD camera (DU-897; Andor Technology) controlled by MetaMorph (Molecular Devices). Emission was filtered 527/27 nm bandpass mirror. Imaging data obtained with either cortical or hippocampal primary cultures did not differ and were therefore pooled together.

### STED Microscopy

Sub-diffraction limited imaging was performed with an Abberior Stedycon instrument as previously described ^13^.

### Image and Data Analysis

Images were analysed using the Fiji edition of ImageJ software ^19^ and the extracted data was processed with Microsoft Excel and GraphPad Prism 10.4.2 release (GraphPad Software, https://www.graphpad.com). Probability distribution of all data sets was assessed in order to apply parametric or non-parametric statistical tests (as stated in figure legends).

## RESULTS

### GABA-B1 receptors localize to neuronal primary cilia in CA1

The ensemble of neuronal GPCRs that are present in the neuronal cilium is not completely described and in addition it is likely to be cell type dependent and dynamic. We therefore performed a series of immunostainings of mouse brain sections looking at ciliary markers and different receptors. We first performed immuno-labeling of mouse brain sections using antibodies against Arl13b, a constitutive ciliary protein, adenylyl cyclase 3 (AC3), a marker of neuronal cilia, and NeuN for detection of neurons. Fig 1A shows a confocal image of the hippocampus documenting the co-localization of Arl13b and AC3 in neuronal cilia. Cilia in non-neuronal, i.e. glial cells, showed stronger Arl13b signal compared to cilia in neurons and lacked AC3. The neuronal primary cilia were also significantly longer than primary cilia of glial cells (Fig 1B). Fig 1C depicts Arl13b combined with an antibody targeting GABA-B1 receptors, revealing the presence of a GABA receptor positive segment, 2-3 µm in length, in a subpopulation of cilia in the CA1 region. Furthermore, cells positive for the glial fibrillary acidic protein did not display ciliary GABA-B1, suggesting that this localization is exclusive of neurons (Fig 1D, E). Some optical sections captured ciliary and dendritic segments that enabled a comparison of GABA-B1 enrichment, and showed that the density of GABA-B1 receptors was higher in primary cilia than in dendrites (Fig 1G). To confirm the sub-ciliary localization, we performed super resolution microscopy using antibodies against AC3 and GABA-B1 receptors. Fig 1F shows a cilium visualized using STED microscopy attesting accumulation of GABA-B1 at the cilium base where it colocalizes with AC3 in the periphery of the cilium, consistent with localization at the ciliary membrane. The averaged longitudinal profile indicates that the length of this sub-ciliary compartment is close to 2.5 μm. To compare the abundance of GABA-B1 in the cilium versus dendrites we analyzed transverse fluorescent profiles taken at the base of the cilium and in dendrites with a comparable width (Fig 1H). The averaged profiles indicate that the ciliary receptor density is equal to or higher than that of adjacent dendrites (Fig 1I).

**Figure 1.**
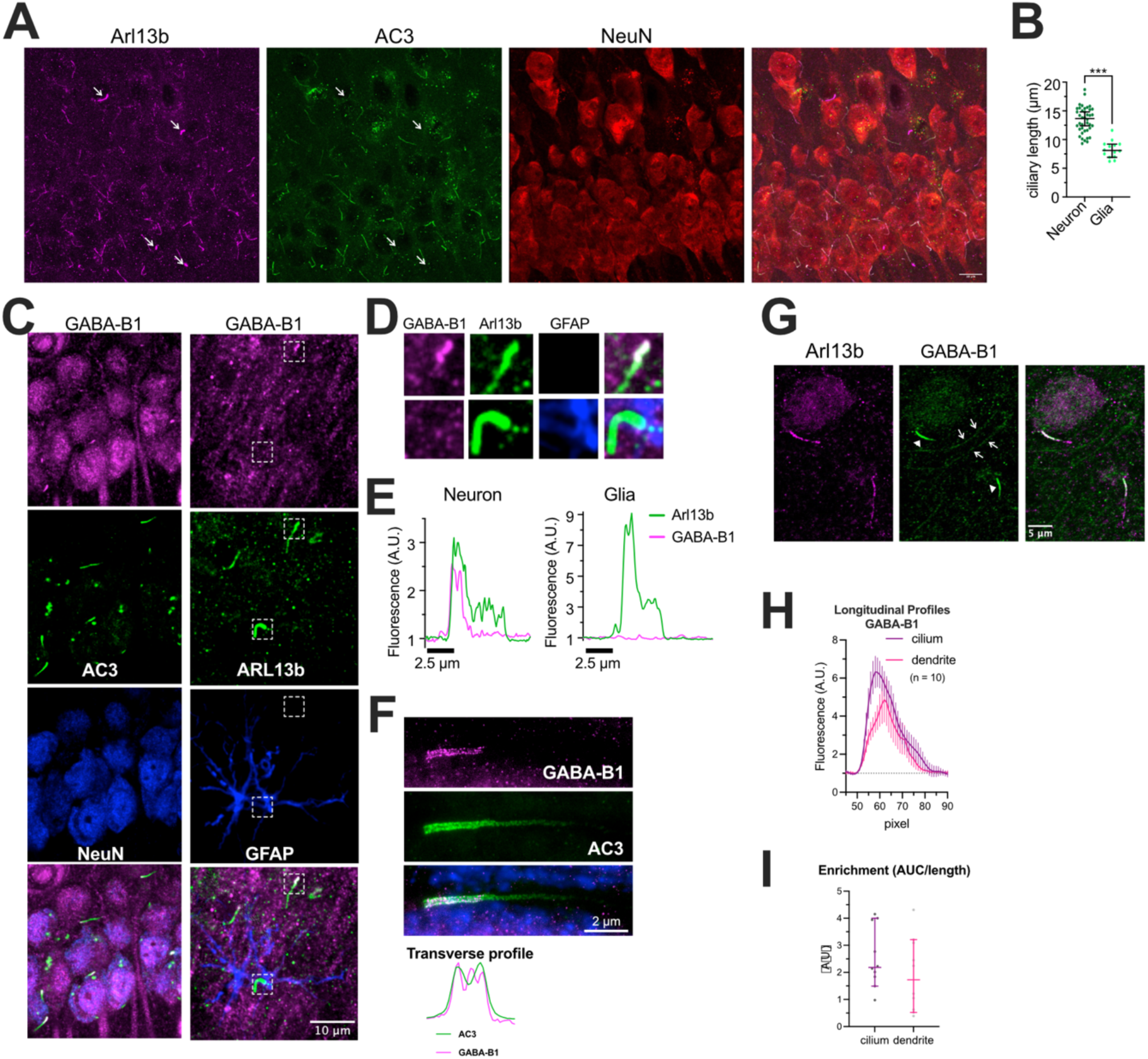
Localization of GABA-B1 receptors to the neuronal primary cilium. **A.** Immunofluorescence images of mouse hippocampal CA1 region showing the distribution of Arl13b (magenta, primary cilium), AC3 (green, neuronal primary cilium) and NeuN (neuronal marker). Arrows show the location of non-neuronal primary cilia. (scale bar 10 µm). **B.** Quantification of cilia length in neuronal and non-neuronal cells in the mouse CA1 (13,38± 0,36µm n=40 neuron and 8,26± 0,40µm n=14 glia, two tailed t-test P<0,0001). **C.** Immunofluorescence images of mouse CA1 showing the distribution of GABA-B1 receptors (magenta), AC3 (green; left column), Arl13b (green; right column) and NeuN (blue; neurons) or GFAP (blue; glial cells). **D.** Magnifications of boxed region in C showing the presence of GABA-B1 receptors on primary cilia in neurons but not in glial cells. Fluorescence intensity along the dashed lines is shown in E below. **E.** Line profiles drawn along a cilium from a neuron (left) and a glial cell (right) showing the presence of GABA-B1 receptors at the cilia base of neuronal primary cilia. **F.** STED microscopy image of a neuronal cilia showing the presence of GABA-B1 receptors at the cilia base, where it colocalized with the cilia membrane marker AC3. The transverse profiles shown below represents the fluorescence intensity along the dashed line. **G.** Immunofluorescence image showing the localization of GABA-B1 receptors to the primary cilium base (arrowheads) and to dendrites (arrows). **H.** GABA-B1 receptor enrichment along the cilia base and a corresponding segment from an adjacent dendrite (n=10). **I.** Relative GABA-B1 receptor enrichment in the primary cilium and dendrites of neurons from the mouse CA1 (n=10, P=0,3930 two tailed t-test).

### Conserved ciliary localization of GABA-B1 receptors across brain regions

Next, we asked whether neurons in other brain regions display GABA-B1 in their cilia as well. We decided to extend our study to the cortex and the striatum, where the majority of cells are the GABAergic medium spiny neurons. In agreement with our observations in the hippocampus, Figure 2A shows that neurons in these two brain regions have cilia that are positive for GABA-B1 while cilia belonging to non-neuronal cells lacked the receptor. Transverse profiles at the base of the cilium and across dendrites in each brain region show again that the receptor density is higher in the cilium (Fig 2B, C) than in adjacent dendrites. Neuronal cilia in the striatum have a length of about 10 μm and a GABA-B1-positive segment of almost 5 μm confined to the cilia base; both significantly longer than their hippocampal and cortical counterparts (Fig 2D). Taken together, these results support the idea of a ciliary GABA-B1 pathway as a neuron specific pathway in glutamatergic as well as GABAergic neurons.

**Figure 2.**
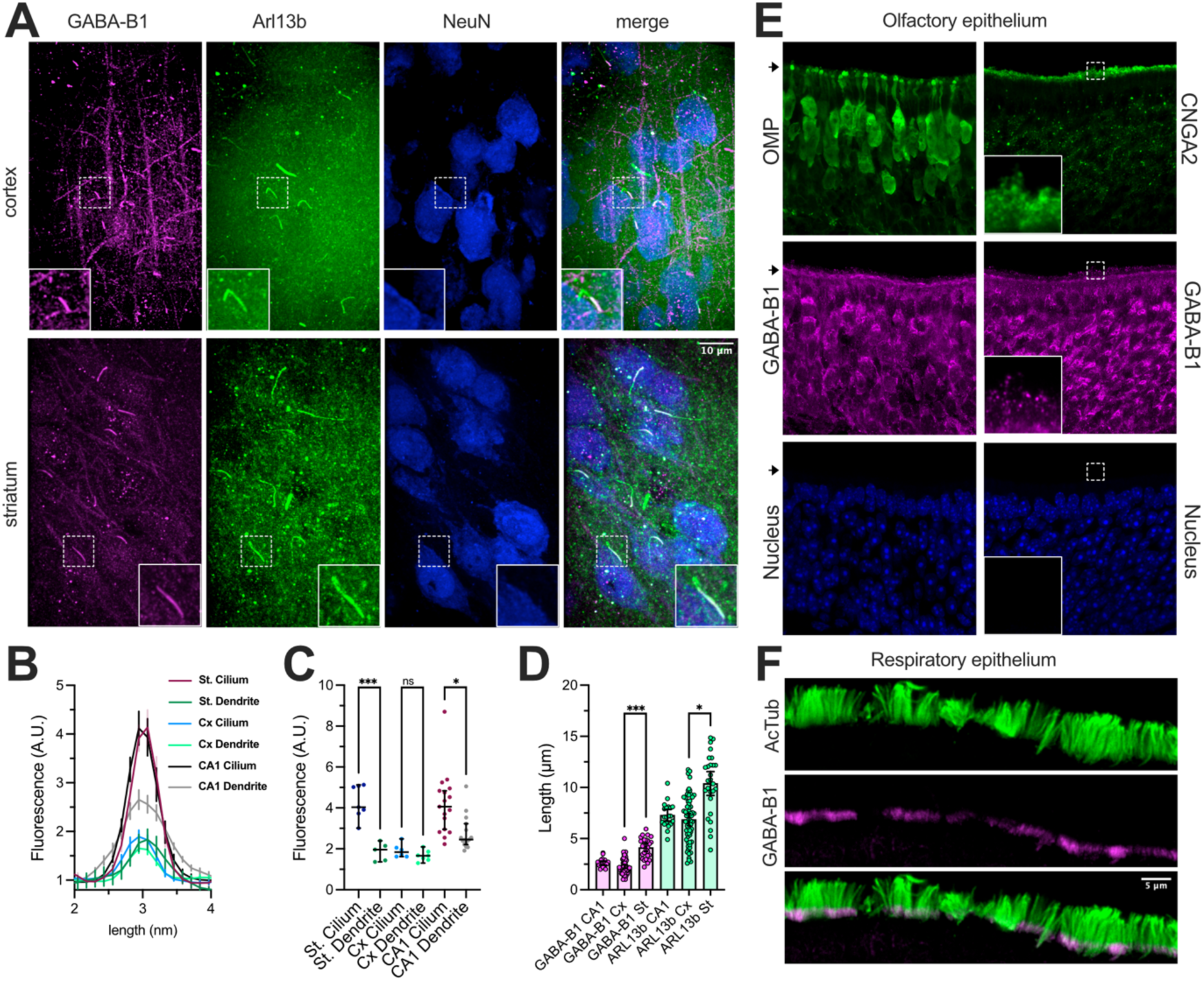
Distribution of GABA-B1 receptors in the mouse brain. **A.** Immunofluorescence images from mouse cortex (top) and Striatum (bottom) showing the distribution of GABA-B1 receptors (magenta), Arl13b (green) and neurons (blue). **B, C.** Quantifications of GABA-B1 immunofluorescence intensity at the cilia base in the cortex, striatum and hippocampus (CA1) in relation to dendrites in the same region (n_cilia/dendrites_: striatum 7/6, cortex 6/7, CA1 17/16). * P=0,0346, *** P=0,0014; Kruskal-Wallis ANOVA followed by multiple comparisons. **D.** Length of primary cilia (green) and the GABA-B1 receptor compartment (magenta) in primary cilia of neurons in the cortex, striatum and hippocampus (striatum_GABA-B1/ARL13b_ n=28, cortex_GABA-B1/ARL13b_ n=64, CA1_GABA-B1/ARL13b_ n=20). *P=0,0189, ***P=0,0001; Kruskal-Wallis ANOVA followed by multiple comparisons. **E.** Immunofluorescence images from mouse olfactory epithelium showing the distribution of GABA-B1 receptors (magenta), OMP (green; left), CNGA2 (green, right) and nuclei (blue). **F.** Immunofluorescence images from mouse respiratory epithelium showing the distribution of GABA-B1 receptors (magenta) and acetylated tubulin (motile cilia; green).

In contrast to most neurons in the central nervous system, olfactory neurons are constantly being renewed and therefore cells present in the olfactory epithelium have different ages. To investigate if this would affect the localization of receptors to the cilia we extended our examination to olfactory neurons. Olfactory neurons were detected by the expression of the olfactory marker protein (OMP, Fig 2E) and lacked enrichment of GABA-B1 in the proximal domain of their cilia. Instead, positive puncta could be seen along the whole length of cilia. Accumulation of the receptor was visible at the junction between apical dendrite and olfactory knob suggesting GABA-B1 receptors are transported to cilia even if not specifically enriched there. In the same preparation motile cilia of the airway epithelium are present and to our surprise a prominent 2 μm long ciliary base segment contained abundant GABA-B1 receptors (Fig. 2F), resembling the segment observed in primary cilia.

Next, we investigated the distribution of other ciliary receptors in the mouse brain by immunofluorescence and confocal microscopy. Melanin-concentrating hormone receptor 1 (MCHR1), known to be present in the cilia of different neurons ^20^, were present in the cilia of neurons residing in CA1 (Fig. 3A, B), and to a lesser extent also in cilia of neurons in the cortex and striatum. Similar distribution was also seen for the hedgehog receptor Patched (Fig. 3B). Interestingly, we were able to detect the presence of prostaglandin E_2_ receptor 4 (EP4) in the cilia of neurons of all three brain areas (Fig. 3B). In contrast to GABA-B1, these three receptors distributed homogenously along the entire cilium (Fig. 3C). We speculated that the distinct localization of GABA-B1 receptors could be due to association with another protein with anisotropic distribution. The inversin compartment has been defined as a set of proteins with a biased proximal localization in cilia of different cell types (Fig. 3D) ^21^. We therefore immunostained mouse brain sections against three proteins belonging to the inversin compartment: ANKS6, NPHP3 and NEK8. While ANKS6 and NPHP3 were absent from neuronal primary cilia, NEK8 was detected in both neuronal and non-neuronal cilia, but in neither cell type was the localization restricted to the proximal segment of the cilium (Fig. 3E). We also failed to detect a distinct inversin compartment in mouse pancreatic islets of Langerhans and clonal insulin secreting cells (Fig. 3E); two preparations where GABA-B1 receptors also show distinct localization to the cilia base ^13^. The inversin compartment therefore seems an unlikely candidate for anchoring GABA-B1 to the base of the cilium in neurons.

**Figure 3.**
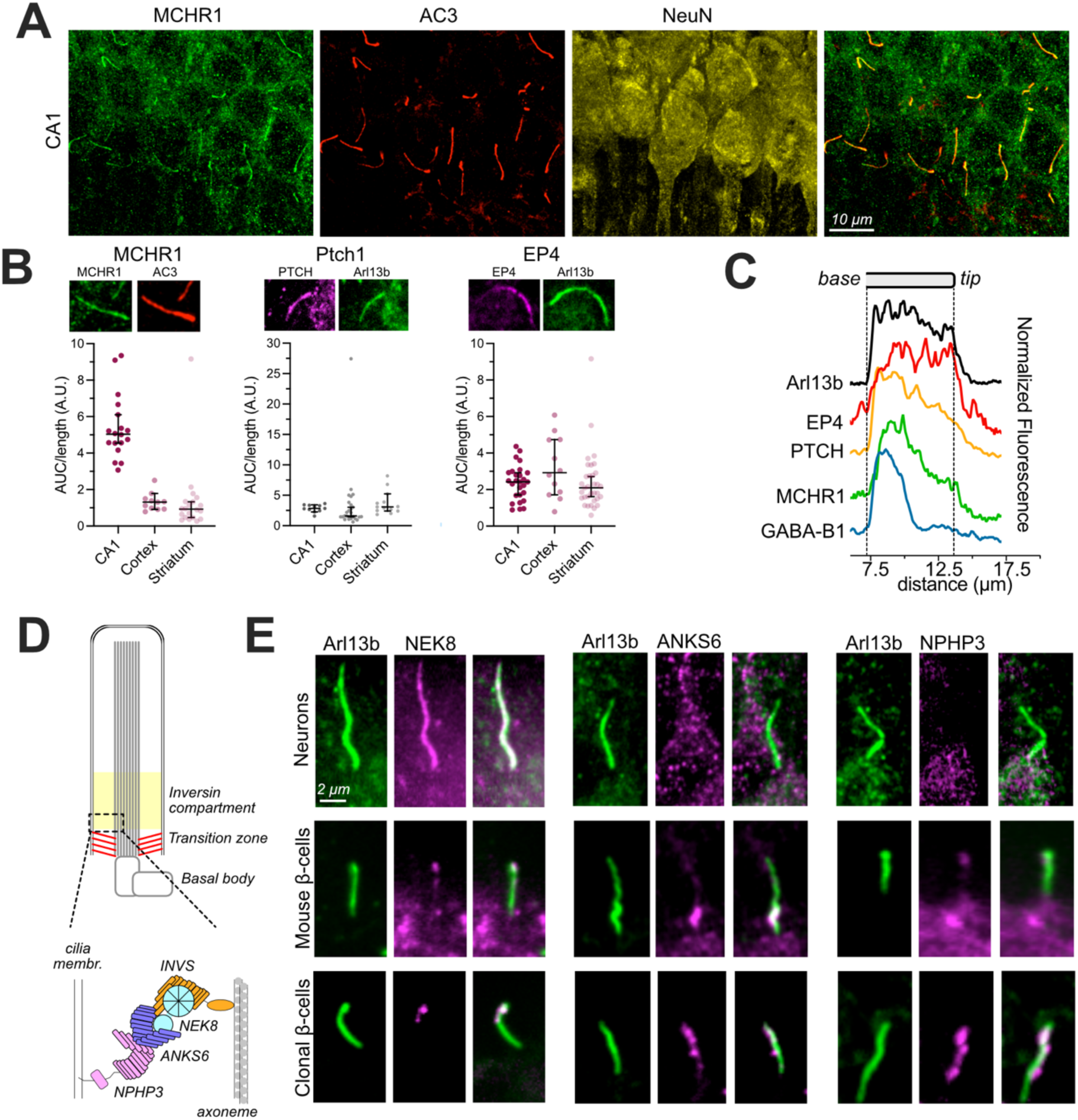
Neuronal primary cilia lack the inversin compartment. **A.** Immunofluorescence images from mouse hippocampus showing the distribution of MCHR1 (green), AC3 (red) and neurons (yellow). **B.** Immunofluorescence images from mouse hippocampus showing the distribution of MCHR1, Ptch1 and EP4 in neuronal primary cilia. Quantification of MCHR1, Ptch1 and EP4 enrichment in neuronal primary cilia relative to the local background in the mouse hippocampus, cortex and striatum are shown below. **C.** Normalized fluorescence intensity profiles along primary cilia of hippocampal neurons showing the distribution (from base to tip) of Arl13b (black), EP4 (red), Ptch (orange), MCHR1 (green) and GABA-B1 (blue). The lines are averages from 13 (EP4), 10 (Ptch), 10 (MCHR1) and 11 (GABA-B1) cilia. **D.** Cartoon showing the proposed localization of the inversin compartment, composed of ANKS6, INVS, NEK8 and NPHP3, distal to the ciliary transition zone. **E.** Distribution of NEK8, ANKS6 and NPHP3 in primary cilia of neurons (top), mouse islet β-cells and clonal MIN6 β-cells.

### Ca^2+^ signaling in neuronal primary cilia

Activation of ciliary GABA-B1 receptors is coupled to local Ca^2+^ influx in insulin secreting cells of the pancreas ^13^. To determine if a similar signal transduction pathway exists in neurons, we employed primary hippocampal cultures that expressed a cilia-localized Ca^2+^ indicator, 5HT_6_-GGECO1 ^16^. Real-time imaging by confocal microscopy revealed spontaneous Ca^2+^ fluctuations in the primary cilium. The events were sparse and required long recording times to be captured, and the Ca^2+^ transients were characterized by quick on and off transitions, a propagation speed close to 5 μm/s and a duration of about 1 minute (Fig. 4A-D). Figure 4E shows an example of decay kinetics at distal and proximal regions of a cilium. When an exponential decay model is fitted to each trace it shows a significantly faster decay at the base and this difference was consistent across events recorded from different cultures (Fig. 4F). This shows that Ca^2+^ clearance from the primary cilium is fastest at the base immediately adjacent to the soma. The Ca^2+^ concentration in neurons fluctuates in response to changes in electrical activity, and since neurons in primary cultures form functional networks, we asked to what extent the ciliary compartment is affected by cytoplasmic Ca^2+^ dynamics. Figure 4G shows representative traces from two neurons with conspicuous and autonomous Ca^2+^ activity; simultaneously the cilium of one of them displays its own activity with little interference from somatic Ca^2+^. To quantify this effect, we looked at the ratio of the Ca^2+^ wave’s peak at the tip and the base of the cilium of the events propagating either tip-to-base (spontaneous flashes) or base-to tip (action potential-derived waves), with the attenuation being stronger for Ca^2+^ moving in the distal direction (Fig. 4H). These results show that neuronal primary cilia are isolated against cytosolic Ca^2+^ concentration changes.

**Figure 4.**
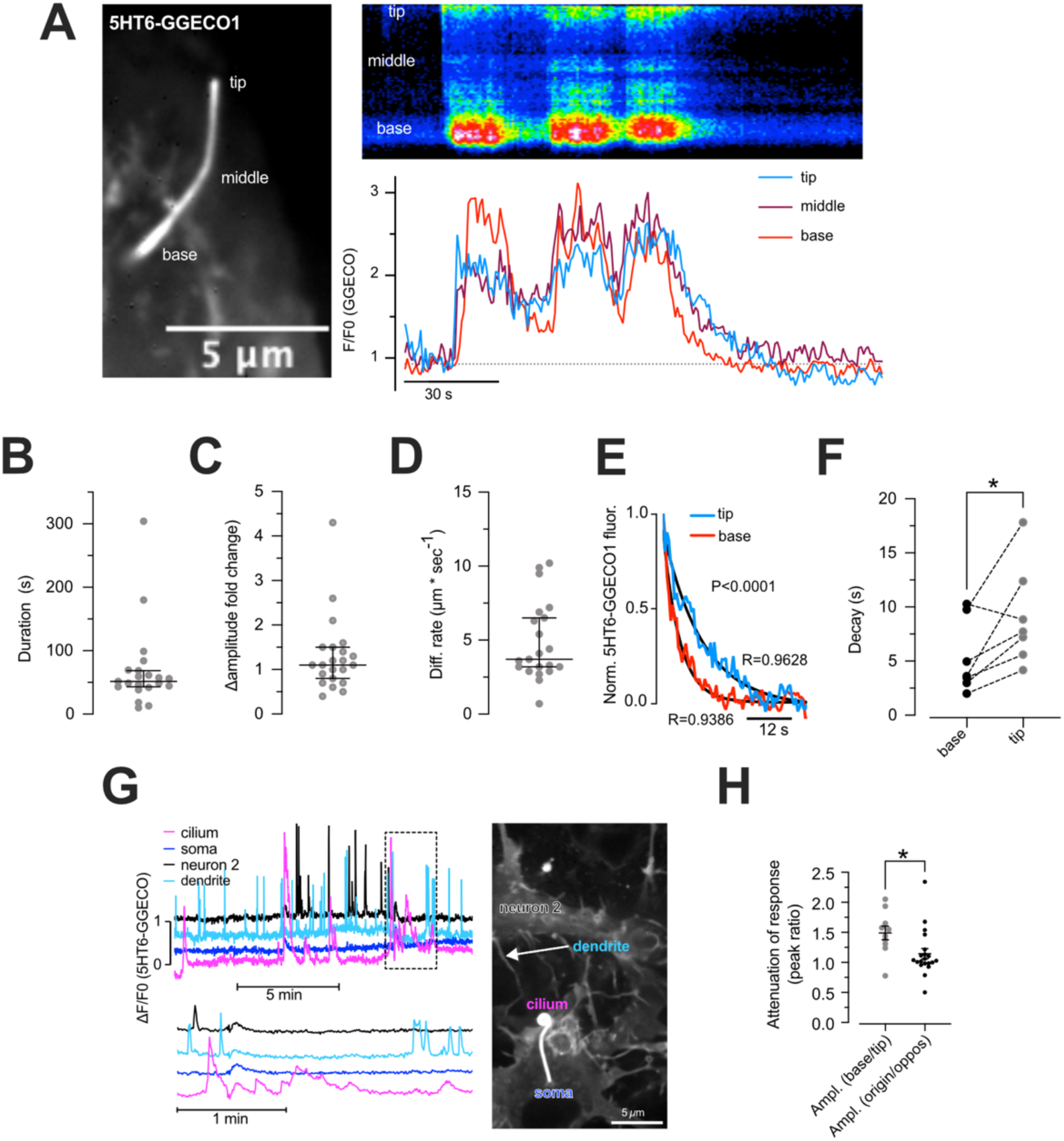
Spontaneous Ca^2+^ signaling in the primary cilium of cultured rat cortical neurons. **A.** Confocal microscopy image of a cultured rat cortical neuron expressing the cilia-localized Ca^2+^ sensor 5HT6-GGECO1. To the right is shown a kymograph from a line drawn along the cilium (top) and 5HT6-GGECO1 fluorescence intensity changes in the cilium base (red), middle (purple) and tip (blue). **B.** Duration of ciliary Ca^2+^ signals in cultured cortical neurons (67,55±13,45 s; n=22). **C.** Amplitude of 5HT6-GGECO1 fluorescence change during spontaneous activity in cultured cortical neurons (1,30±0,17 A.U.; n=23). **D.** Ca^2+^ diffusion rate inside primary cilia of cultured cortical neurons (4,79±0,56 μm*s^-1^; n=22). **E.** Ca^2+^ decay kinetics at the cilium base and tip (from panel A). **F.** Ca^2+^ decay half-life in the base (solid) and tip (open) segments of primary cilia in cultured cortical neurons (n=7; P=0,0312; two tailed Wilcoxon matched-pairs rank test). **G.** Recordings of spontaneous Ca^2+^ activity in cultured rat neurons expressing 5HT6-GGECO1. The traces show 5HT6-GGECO1 fluorescence changes in the primary cilium (magenta), dendrite (light blue), soma (dark blue) and an adjacent neuron (black). Boxed region is shown below on an expanded time-axis. **H.** Action potential derived ciliary wave attenuation (base/tip max amplitude ratio, n=10) is more potent than ciliary transduction calcium attenuation (site of origin/opposite region max amplitude ratio, n=21; P=0,0101 two tailed Mann-Whitney test).

### GABA-induced Ca^2+^ signaling in neuronal primary cilia

Having demonstrated the presence of GABA-B1 receptors and local Ca^2+^ signaling in the primary cilium of neurons, we next asked whether the receptor could be, at least in part, responsible for transducing GABAergic signals into ciliary Ca^2+^ transients. We first confirmed the presence of ciliary GABA-B1 receptors in both cultured naïve neurons and neurons expressing the ciliary Ca^2+^ reporter (Fig. 5A). As expected, expression of the Ca^2+^ indicator induced a considerable lengthening of the cilia, though the restricted proximal localization of the GABA-B1 receptor was maintained (Fig. 5B).

**Figure 5.**
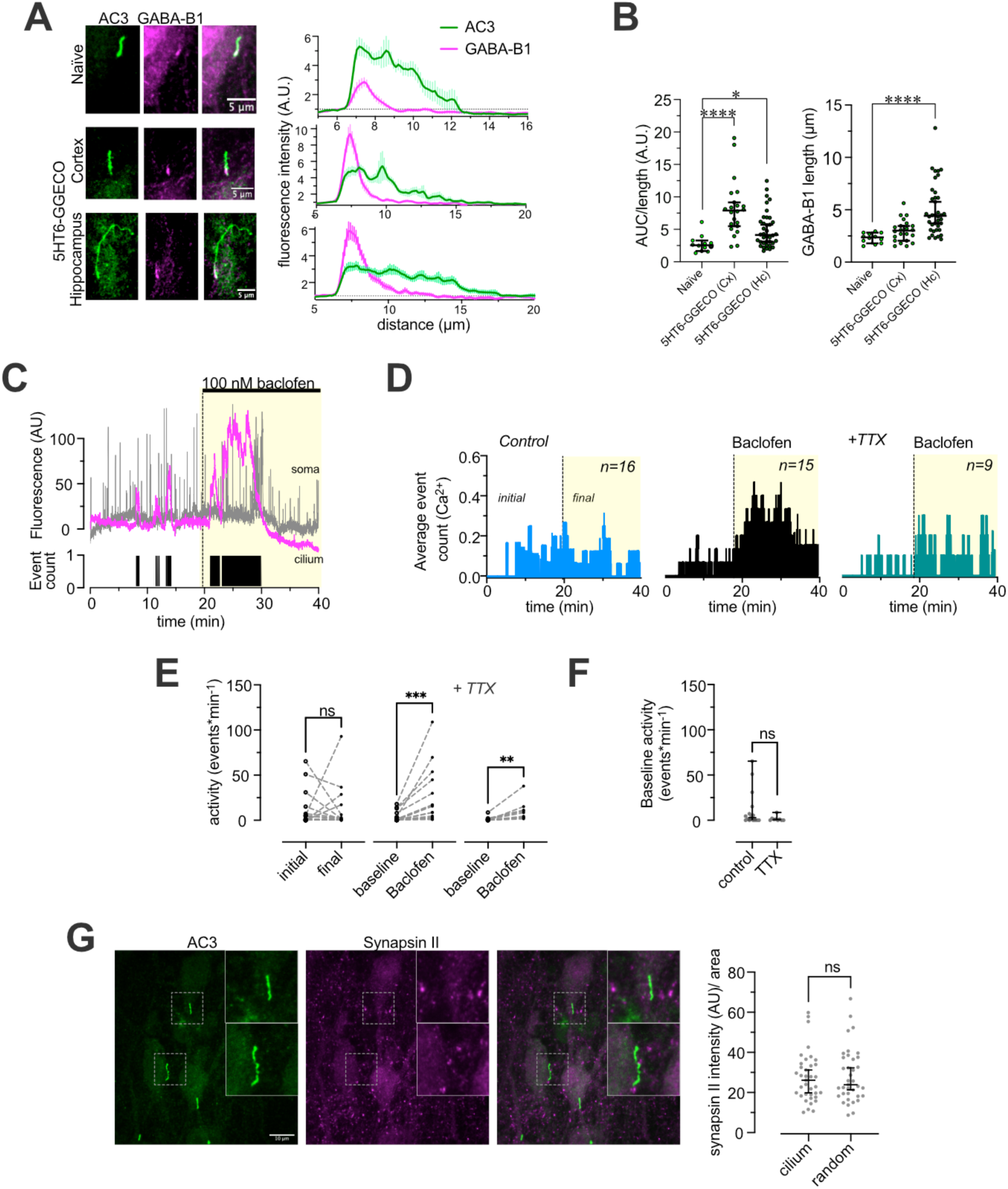
GABA-B1 receptor activation triggers Ca^2+^ signaling in the primary cilia of cultured rat cortical neurons. **A.** Immunofluorescence images from naïve or 5HT6-GGECO1-expressing cultured mouse neurons (from cortex and CA1) showing the distribution of GABA-B1 receptors (magenta) in primary cilia positive for AC3 (green). Intensity profiles from lines drawn along the cilia are shown to the right and further show the confinement of GABA-B1 receptors to the cilia base of cultured neurons. **B.** Quantification of GABA-B1 receptor enrichment at the cilia base in naïve cultured neurons or cultured cortical and hippocampal neurons expressing 5HT6-GGECO1 (naïve 2,76±0,37; Cx 8,18±0,91; Hipp 4,94±0,44 arbitrary units). GABA-B1 positive segment length was measured for each group and found significantly different (naïve, Cx and Hipp, in µm: 2,78±0,37; 8,18±0,91; 5,03±0,39; mean±SEM; Kruskal-Wallis ANOVA followed by multiple comparisons, naïve n=12, Cx n=22, Hipp n=38, for **A** & **B).** **C.** Simultaneous recording of cytoplasmic (grey) and ciliary (pink) calcium activities from a neuron before and after stimulation with 100nM baclofen. Activation of metabotropic GABA receptors was without effect on the frequency of somatic events while ciliary signaling was increased by the agonist. Positive deviations from baseline were computed over time to produce ciliary activity count traces (black, lower panel). **D.** Averaged activity counts corresponding to neurons in control (blue) baclofen (100nM, black) and TTX (1 µM) + baclofen (100nM, grey) showing an increase of activity after stimulation with baclofen. **E.** Quantification of activity changes was performed in a pair wise fashion, comparing the averaged activity during initial vs final period (control) and baseline vs baclofen (either in the absence of presence of TTX). Time lacked an effect on activity (control P=0,4332) while baclofen stimulation led to a significant increase in both - /+ TTX conditions (respectively P=0,0002 and P=0,0039; two tailed Wilcoxon matched-pairs signed rank test) **F.** To test whether abolition of action potentials had an impact on spontaneous ciliary activity, recordings in control and TTX (initial 20 minutes both groups) were compared. Statistical analysis showed no difference (P=0,1468 Mann-Whitney two tailed t-test). **G.** Immunofluorescence images showing the distribution of primary cilia (AC3; green) and synapses (Synapsin II; magenta) in cultured cortical neurons. Boxed areas are magnified to the right. Scatter plot to the left shows Synapsin II fluorescence intensity at primary cilia and random locations within the same sample (P=0,3696 Wilcoxon matched pairs signed rank test).

To test the functionality of the ciliary GABA pathway, we activated metabotropic GABA receptors with a low dose of the agonist baclofen (100 nM). The addition of baclofen produced an increase in ciliary Ca^2+^ activity and quantification of the aggregate activity per minute for each recording before and after treatment showed that baclofen increased the activity while there was no time effect in the control experiments (Fig. 5C-E). Finally, to investigate the source of GABA driving ciliary signaling we included 1 μM TTX in the extracellular buffer to block sodium channels. In the absence of action potential evoked synaptic transmission, ciliary activity was not abolished and baclofen could still enhance Ca^2+^ signaling in the primary cilium, although the response magnitude was slightly attenuated (Fig. 5D-F). Consistent with this, we did not observe any evidence of proximity or bias in localization of neuronal cilia in relation to synapses positive for the presynaptic marker Synapsin II (Fig. 5G).

## DISCUSSION

Neurons are highly compartmentalized cells in which individual synapses constitute the computational units of neural networks ^22^. Here we show that the neuronal primary cilium is an organelle participating in cell-to-cell communication, and that GABA, through cilia-localized GABA-B1 receptors, is one of the messenger molecules.

GABA is the major inhibitory neurotransmitter in the brain and activates either ionotropic or metabotropic receptors. The finding that G-protein-coupled GABA-B1 receptors can drive ciliary Ca^2+^ transients is perplexing but recapitulates our recent findings in pancreatic β-cells ^13^. In addition, we show by immunostaining that the presence of GABA-B1 receptors in the cilium is conserved across different brain regions, suggesting that it is a common feature of most neurons in the adult brain. The absence of the receptor from cilia of non-neuronal cells, including GFAP-positive astrocytes, further highlights that this signalling pathway is restricted to neurons. The GABA-B1 receptor is confined to a sub-ciliary compartment; a ∼ 2,5 μm long segment at the base. This proximal domain appears to be invariant across different cell types that include neurons, β-cells and motile cilia of airways epithelial cells (but not olfactory neurons and glial cells). It is a peculiar distribution pattern not adopted by any other known ciliary receptor, and the reasons behind this characteristic are elusive. The size and localization of the compartment is reminiscent of the inversin compartment, which is a structure composed of INVS, ANKS6, NPHP3 and NEK8 that is involved in ciliary signaling, in particular during development ^21^. Examination of the inversin compartment components in adult mouse brain sections revealed that only NEK8 was present in the cilium, but its distribution was not confined to the proximal part. Similar observations were made in mouse endocrine islets of Langerhans, thus questioning the inversin compartment as a conserved feature of primary cilia, at least in cells with little or no proliferation. In any case, the inversin compartment is not responsible for anchoring GABA-B1 receptors to the cilia base. How the GABA-B1 receptor gets into the cilium is also an open question, since it lacks typical targeting sequences found in GPCRs ^23^. Metabotropic GABA-B1 receptors heterodimerizes with GABA-B2 receptors, and this interaction is required for both plasma membrane targeting and G-protein-coupling ^24^ ^25^, but whether this interaction is also required for cilia targeting is not known. While GABA-B1 receptors are ubiquitously expressed in human tissue, GABA-B2 receptor expression is restricted to the central nervous system (Human protein atlas). This difference in expression pattern indicates that GABA-B1 receptors can function independent of GABA-B2 receptors, and one possibility is that this function involves the primary cilium.

Ca^2+^ imaging in neuronal cultures showed that the cilium of neurons was isolated from cytoplasmic Ca^2+^ interference. Cytoplasmic Ca^2+^ increases, caused by spontaneous neuronal activity, were prevented from propagating into the more distal sections of the cilium, and this is likely achieved by efficient clearance in the proximal domain ^13^. At the same time, we observed spontaneous Ca^2+^ signaling in the primary cilia of neurons that was distinct from cytosolic Ca^2+^ signaling and also did not propagate beyond the cilia base. These primary cultures were deprived of non-neuronal cells due to omission of serum from medium, indicating that neurotransmitters utilized by neurons, GABA among them, may be the trigger of these Ca^2+^ signals. These events were rare, but we expect that *in vivo* preparations, where both cytoarchitecture and complexity of the signaling landscape are preserved, should display significantly more activity. Inhibition of action potential-driven synaptic release with TTX was without effect on spontaneous ciliary Ca^2+^ signaling, indicating that the source of these events is non-synaptic. Alternative sources include extracellular ligands released in a non-synaptic fashion ^26^ ^27^ and intra-ciliary ligands, such as cAMP or cGMP ^28^.

Recent studies have shown that neuronal cilia are positioned to sense signals emanating from different origins. A functional axo-ciliary connection has been described in CA1, while other structural studies document the integration of cilia in traditional synapses or their close proximity to them ^8^ ^7^ ^5^. Our observation did not show any evidence of directs synapses involving neuronal cilia, although we do not rule out that such may exist in certain neuronal sub-populations. Studies of intact tissue will be required to establish the origin of the signals that activate second messenger dynamics in the cilium. Additionally, the cilium is well suited for increasing signal transduction sensitivity by providing a reduced two-dimensional diffusion space concomitantly to the high concentration of ciliary membrane proteins, and therefore the organelle could in principle be responsive to subtle changes in e.g. volume transmission. Neuronal cilia have been associated with different cognitive functions including food intake ^29^ and circadian rhythm ^30^, as well as in pathological states such as Alzheimer’s disease ^31^ and Parkinson’s disease ^32^. Interestingly, both Alzheimer’s disease and Parkinson’s disease are also characterized by aberrant GABA signaling ^33,34^, and understanding whether ciliary GABA signaling may be involved in disease pathogenesis is an important future research aim.

## ACKNOWLEDGEMENTS

This work was funded by grants from the Swedish research council and the Swedish brain foundation (Hjärnfonden) (to OI-H) and from OE&E Johanssons foundation (to GS). We also acknowledge the staff at the local microscopy core facility BioVis for help with STED microscopy and to members of the OI-H and ML labs for input on the work. None of the authors have any competing interests to declare.

